# The assessment of cortical hemodynamic responses induced by tubuloglomerular feedback using in vivo imaging

**DOI:** 10.1101/2022.09.19.508507

**Authors:** Blaire Lee, Dmitry D. Postnov, Charlotte M. Sørensen, Olga Sosnovtseva

**Affiliations:** Department of Biomedical Sciences, Copenhagen University, Blegdamsvej 3, 2200 Copenhagen N, Denmark; Department of Clinical Medicine, Aarhus University, Aarhus, Denmark

## Abstract

The tubuloglomerular feedback (TGF) mechanism is crucial in modulating renal hemodynamics and glomerular filtration rate in individual nephrons. Our study aimed to evaluate the TGF-induced vascular responses by comparing the effects of two transport inhibitors with different sites and mechanisms of action. We assessed cortical hemodynamics with high-resolution laser speckle contrast imaging, which enabled the evaluation of blood flow individual micro-vessels and analysis of their dynamical patterns in the time-frequency domain. We demonstrated that a systemic administration of a loop diuretic abolishes TGF-mediated hemodynamic responses. Furthermore, we showed that the local microcirculatory blood flow decreased, and the TGF reset in response to reduced proximal reabsorption elicited by systemic administration of a sodium-glucose co-transporter 2 inhibitor, phlorizin.

## 1 Introduction

Despite the constant fluctuation in systemic blood pressure, the kidney maintains a stable filtration rate due to renal autoregulation [1] which takes place in individual nephrons, or structurally discernible filtration units of the kidney. The two prominent components of renal autoregulation are the myogenic response (MR) and the tubuloglomerular feedback (TGF) mechanism. While both components target the resistance of afferent arterioles, they occur in response to distinct triggers. MR is the constriction of a vessel due to an increase in transmural pressure. TGF correlates to the amount of chloride passing into the distal tubule. When the macula densa detects an increase in the tubular fluid Cl^−^ concentration through the Na-K-2Cl co-transporters, it signals the afferent arteriole to constrict [2, 3, 4, 5]. The effect is reduced blood flow and hydrostatic pressure into the glomerulus. Managing glomerular capillary pressure is pertinent for all kidney models to sustain function and prevent deterioration.

Since the concentration of tubular sodium chloride modulates TGF, pharmacological modulation of the detection and the reabsorption of the ions by inhibiting various Na^−^ transporters is a viable therapeutic target. Furosemide is a loop diuretic for treating patients with heart failure and hypertension. It inhibits Na-K-2Cl co-transporters (NKCC2) expressed in Henle’s thick ascending loop and the macula densa for active reabsorption and detection of chloride, respectively [6, 7]. In parallel, it is well-known that inhibiting the detection of tubular chloride at the macula densa using loop diuretics suppresses TGF in a single nephron [2], but this has not been observed in a population of nephrons.

The kidney plays an integral role in glucose homeostasis partly through glucose reabsorption. Sodium-glucose co-transporter 2 (SGLT2) expressed in the luminal membrane of the proximal convoluted tubules is responsible for 97% of the glucose reabsorption through a sodium-dependent active transport [8]. Phlorizin, an SGLT2 inhibitor, lowers the blood glucose level, induces glucosuria, and promotes weight loss in diabetic patients by blocking the sodium-glucose reabsorption [9, 10]. Recent findings suggest that inhibiting SLGT2 can reduce glomerular hyperfiltration restoring TGF in a compromised kidney [11], but the direct effect of SGLT2 inhibition on nephron blood flow and TGF remains poorly understood.

High-resolution laser speckle contrast imaging (LSCI) can measure microvascular hemodynamics in a population of nephrons in real-time [12, 13]. TGF modulates the afferent arteriolar resistance at around 0.033 Hz resulting in oscillatory blood flow. LSCI can capture this dynamic at a high spatio-temporal resolution. Thus, high-resolution LSCI is an apt modality for investigating transient perturbations to TGF oscillations and the resulting renal hemodynamics.

We acutely administered systemic infusions of furosemide (NKCC2 inhibitor) or phlorizin (SGLT2 inhibitor) in anesthetized rats. We assessed the TGF-induced hemodynamic changes in a population of nephrons *in vivo* with a high-resolution LSCI. To characterize the TGF, we implemented the time-frequency superlet analysis, an improved method for analyzing bio-signals [14]. We then extracted new analytical metrics to quantify the characteristics of TGF oscillations. Finally, we evaluated the effects of furosemide and phlorizin on cortical hemodynamic responses mediated by TGF.

## 2 Materials and Methods

### 2.1 Surgical preparation and experimental protocol

Danish National Animal Experiment Inspectorate approved all rodent experiments. Male normotensive Sprague Dawleys (N=13, RjHan:SD, Janvier) weighing 330±23 g were used. The animals were fed *ad libitum* and housed in a 12-hour light/dark cycle. Detailed animal preparation can be found in Refs [15, 13]. Here we provide an abridged version. All experiments were performed under sevoflurane anesthesia (Sevorane, Abbvie), 8 % at induction, and maintained at around 1.5 %. The body temperature was maintained at 37°C on a servo-controlled heating table. The animal was laid ventral side up, and a mask delivering anesthesia covered the nose and the mouth during the surgery. A small horizontal incision was made between the cervical and pectoral region to expose and isolate the trachea, the left jugular, and the right common carotid. Two polyethylene tubes (PP10, Smith Medicals) were inserted into the left jugular vein for infusion of saline (0.5 % NaCl) and muscle relaxant (0.5 mg/ml) (Nimbex, Apsen) at 20 *μ*l per minute via syringe pumps (AL-1000, World Precision Instruments). A slightly larger polyethylene tube (PP50, Smith Medicals) was placed into the right carotid for continuous blood pressure monitoring. The animal was ventilated through tracheotomy with a volume-controlled respirator (7025, Ugo Basile) at 7 ml per stroke (60 strokes/minute). Laparotomy was performed to expose and stabilize the kidney with a custom 3D printed well. 1.5 % (w/v) agarose solution was poured over the kidney, and a glass coverslip was gently placed on top of the kidney to prevent a parched surface. A myograph wire(40 *μ*m in diameter) bent at an acute angle was then placed on top of the coverslip to track visceral and breathing motion for image registration. The left ureter was cannulated with a polyethylene tube (PP10 connected to PP50, Smith Medicals) for free urine flow and sample collection. A renal flow probe (1PRB, connected to T420 Transonic) was fastened around the renal artery for one animal to measure the standard renal blood flow (4.86 mL/min). Each experiment was considered successful for animals with a stable blood pressure of around 100-130 mmHg.

The animals were separated into two groups: Animals receiving furosemide (n=5) and animals receiving phlorizin (n=7). First, baseline saline infusion and image recording took place for 30 minutes. Then, the saline syringe connecting to the left jugular was replaced with either Phlorizin (50mg/kg, 274313, Sigma-Aldrich) or furosemide (10mg/kg) to induce glucosuria or diuresis, respectively [16]. A minimum of 30 minutes of acclimation period allowed the administered substance to reach its peak systemic concentration. Finally, thirty minutes of post-infusion recording took place while continuing the infusion to sustain bioavailability and saturate the TGF response. See the experimental overview in Fig. 1.

**Figure 1:**
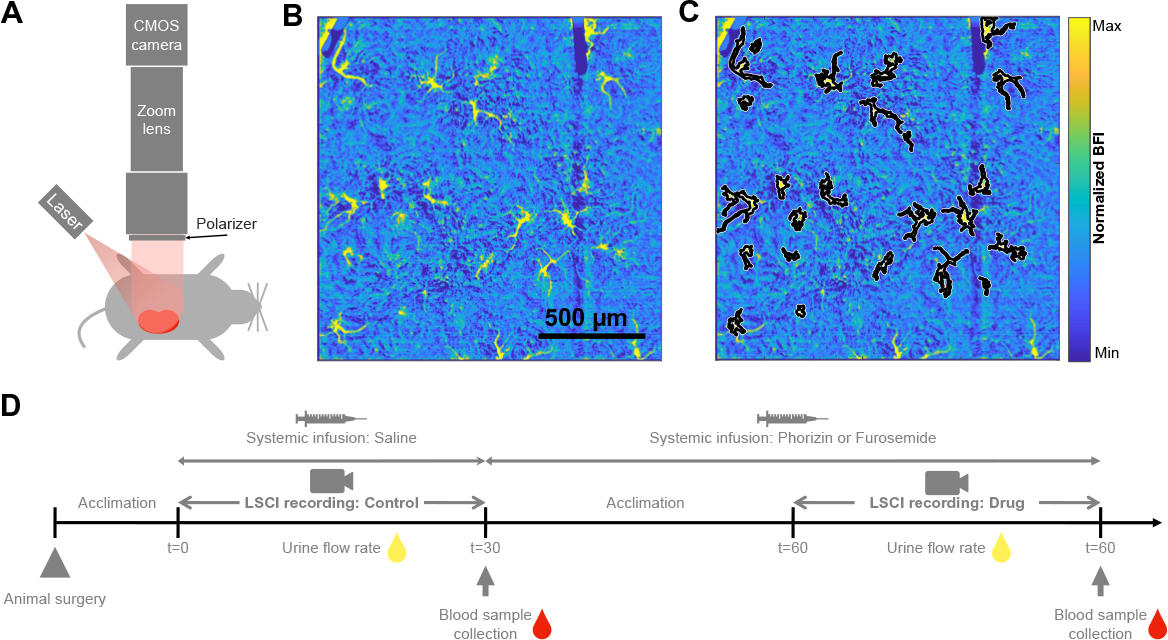
Experimental overview assessing the influence of loop diuretic (Furosemide) and SGLT2 inhibitor (Phlorizin) on renal hemodynamics. A) Imaging setup for laser speckle contrast imaging for real-time high-resolution tissue blood flow measurements. B) Time-averaged blood flow index map acquired with the LSCI. C) An example of segmented micro-vessels. D) Timeline for the whole experiment.

### 2.2 Imaging setup

We used a high-resolution LSCI [12] to record the changes in renal blood flow with the infusion of Phlorizin or Furosemide. Here we briefly describe the imaging setup. A CMOS sensor camera (Basler acA2048-90umNIR, 5.5 *μ*m pixel size, 8bit mode) was vertically attached to a video zoom lens with 4.5 x magnification (VZM 450, Edmund optics). A volume holographic grating stabilized diode (LP7850-SAV50, 785 nm Thorlabs) was used to illuminate the tissue surface and passed through a linear polarizing filter before reaching the photo-sensors to reduce recorded specular reflections and first-order scattering events. Finally, the apparatus was mounted to a height-adjustable z-stage plate to enable vertical translation. Thirty minutes of raw speckle images were recorded at 50 Hz with 5 msec exposure time for the control and the post-drug infusion period. We eliminated all ambient light and kept all settings and system configurations consistent across experiments.

### 2.3 Data analysis

Post-acquisition data analysis approaches are described in Refs [12, 13]. Here we provide a summary of each step.

#### 2.3.1 Image registration and speckle contrast analysis

We aligned the images to the first frame to eliminate the motion artifact from breathing in the x-y plane. A thin myograph wire was placed on top of the glass cover over the kidney as a reference mark during the experiment. The raw frames were converted to binary images, with the myograph wire and the surrounding tissue as the foreground and the background. We then calculated the geometrical transformation for each frame compared to the first frame and aligned the images based on transformation metrics.

We implemented temporal contrast analysis to preserve high spatial resolution to improve image quality and segmentation performance. First, temporal laser speckle contrast was calculated from raw data sets by 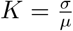. σ is the standard deviation, and *μ* is the mean of the selected temporal kernel (25 frames). Following temporal contrast analysis, the data was sub-sampled from 50 Hz to 1 Hz and converted from the contrast images to blood flow index images by 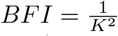.

A mean spatial blood flow map was created by averaging the frames recorded over the time frame of interest: control and drug. Semiautomatic segmentation was applied to high zoom data to segment individual vessels from the surrounding renal tissue according to the approach described in Ref [13, 12].

### 2.3.2 Time-frequency analysis

TGF is an oscillatory signal that operates within a range of frequencies (0.018 and 0.033 Hz), and it can vary over time like many other biological oscillations [17]. Therefore, time-frequency superlet analysis was implemented to the extracted flow time series to reveal TGF oscillations[14]. While there are many methods to analyze frequency components of bio-signals, superlet was chosen for its exceptional performance in resolving the frequency of a signal that changes over time, a primary objective in detecting the changes in TGF frequencies over an observation period [14].

We extracted three metrics from the blood flow time series and the power spectrum. These metrics were compared between the control and the drug period.

- Metric 1 (BFI): Mean blood flow index was calculated by averaging the blood flow index time series over the control and the drug period.
- Metric 2 (Sigma). The standard deviation of TGF oscillations. Blood flow index time series were passed through a band-pass filter of frequency between 0.015 Hz and 0.04 Hz so that signals only contained the frequency range of interest. Then we calculated the standard deviation of filtered signals. We use this metric to measure the amplitude of TGF oscillations.
- Metric 3 (AUC). The area under the curve of the power spectrum within the TGF frequency. First, we performed trapezoidal numerical integration between 0.015 Hz and 0.04 Hz of the individual power spectrum to find the area under the curve within the TGF frequency band. Then we drew a median line between 0.04 and 0.05 Hz for each power spectrum and calculated the area below the line within the TGF frequency band. Then the area below the median line was subtracted from the area under the curve. We use this metric to measure the significance of TGF band frequencies.

### 2.3.3 Statistical analysis

Paired sample t-test was used to compare the urine samples collected before and after administering either furosemide or phlorizin. In addition, ANOVA for the linear mixed-effects model was used to compare the TGF metrics between control and their respective furosemide or phlorizin data, adjusting for between-subject variability. Results with p-values less than 0.001 were declared significant. Bar plots are expressed as mean ± SE.

## 3 Results

### 3.1 Characterizing the TGF-induced hemodynamics

To accurately capture the transient dynamics of TGF, we took an exemplary micro-vessel blood flow time series obtained from a random experiment. We decomposed the signal from the time domain to frequency-power components using time-frequency superlet analysis shown in Fig. 2. It has been demonstrated that the superlet provides a superior time-frequency resolution compared to the Fourier and the Wavelet transform. The time-averaged power spectrum shows a sharp and narrow peak near 0.3 Hz, demonstrating that the superlet analysis can sharply localize the TGF frequency as shown in Fig. 2E. Fig. 2F shows the TGF signal over time for the control period, which corresponds to the oscillation shown in Fig. 2C. The line at 0.3 Hz disappears on the right panel of Fig. 2F because the TGF oscillation no longer exists, demonstrated by the noisy signal in Fig. 2D.

**Figure 2:**
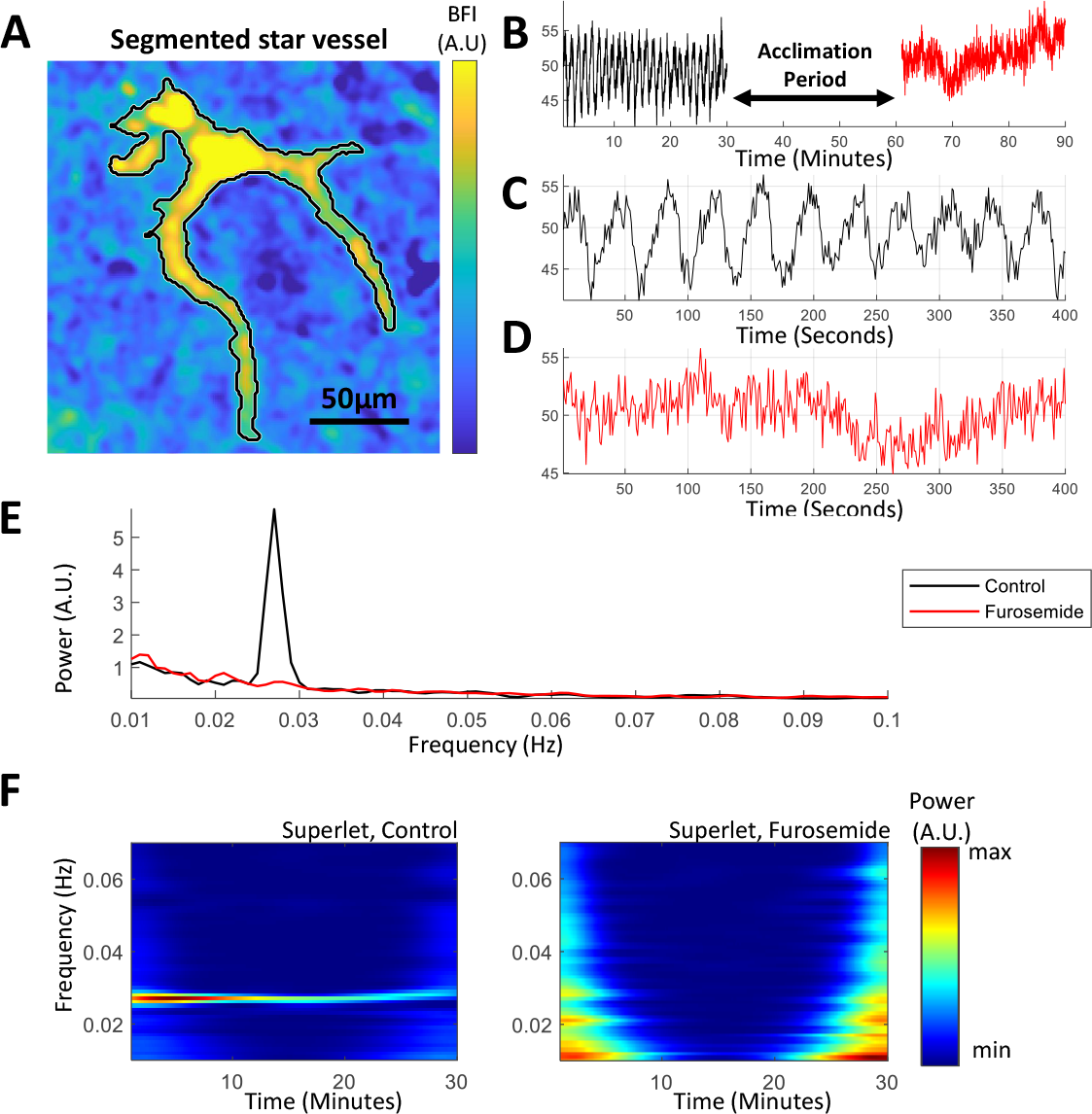
Illustration of the analysis from segmentation to time-frequency spectrogram. A) Mean blood flow index image of a segmented micro-vessel. B) Exemplary blood flow index time series from a segmented vessel. C and D) A closer look at B reveals the TGF oscillations during the control period (black) that disappear with furosemide infusion (red). E) Power spectrum shows a prominent peak at around 0.03 Hz associated with TGF F) Spectrogram of superlet analysis for control period (left) shows a sharp localization of the TGF frequency around 0.03 Hz for 30 minutes that disappears completely with furosemide infusion (right).

With the implementation of superlet analysis, we explored how nephron hemodynamics can vary from vessel to vessel. Although TGF is known to be a persistent mechanism in the nephrons, we found three distinct types of hemodynamic behaviors in the power spectrum, shown in Fig. 3. In some nephrons, TGF operates at a single frequency stable over the observation period (Fig. 3A). In other nephrons, we did not see any TGF peaks in the power spectrum (Fig. 3B). Although this does not mean the TGF mechanism does not exist, it may be indistinguishable from other dynamics and noise. In many cases, we found that nephrons can have dynamic TGF frequencies over time, exhibiting multiple peaks between 0.015 Hz and 0.04 Hz (Fig. 3C). For example, the time-frequency spectrogram in Fig. 3F shows that the TGF frequency slowly migrates from 0.02 Hz to 0.025 Hz over the observation period of 900 seconds.

**Figure 3:**
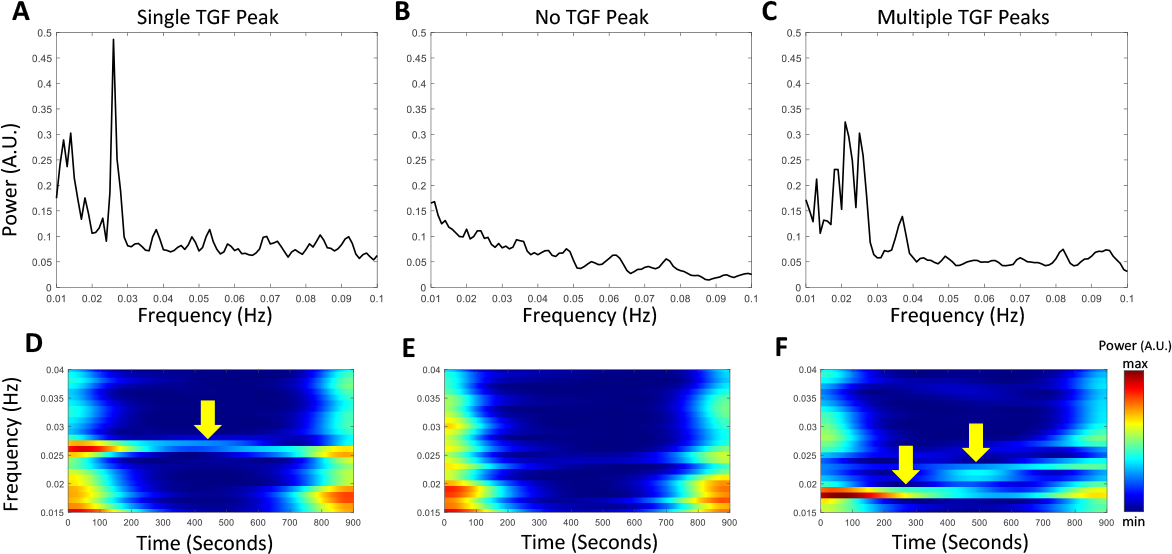
Three types of TGF-induced responses in different vessels. The top panel represents time-averaged power spectra: One peak around 0.03 Hz corresponds to a single frequency of TGF oscillation (A); a TGF peak cannot be detected (B); Multiple peaks within the narrow frequency band correspond to TGF oscillations with different frequencies (C). The bottom panel represents the time-frequency spectrogram matching to power spectra on A-C: A stable TGF frequency over the observation time (D) No observable TGF frequency (E); Multiple TGF frequencies within the observation time (F).

### 3.2 Effect of acute Na-K-2Cl co-transporter blockade on TGF dynamics

Furosemide, a loop diuretic that blocks Na-K-2Cl-co-transporters, is known to cause diuresis and increase the excretion of sodium, chloride, and other ions. In animals receiving the systemic infusion of furosemide, urine flow increased from 12.6±2.95 to 63.0±3.63 *μ*L/min (*P* < 0.01). The urinary sodium excretion increased from 0.38±0.29 to 5.98±0.57 *μ*Eq/min (*P* < 0.01). The blood pressure remained relatively unchanged from 112.16 ±3.21 to 108.49 ±2.21 mmHg.

With the systemic infusion of furosemide, we aimed to abolish TGF activity and observe the hemodynamic changes. Fig. 4 shows that for animals with TGF oscillations during the control period (animals 1-3), it becomes suppressed after administration of furosemide, demonstrated by the lack of vertical striations (second column) and the absence of red peaks in averaged power spectra (third column). In animals 4 and 5, there was a lack of TGF oscillations during the control period, demonstrated by the lack of color contrast in the striations in the carpet plot and no black peaks in their respective power spectra shown in the third column of Fig. 4B.

**Figure 4:**
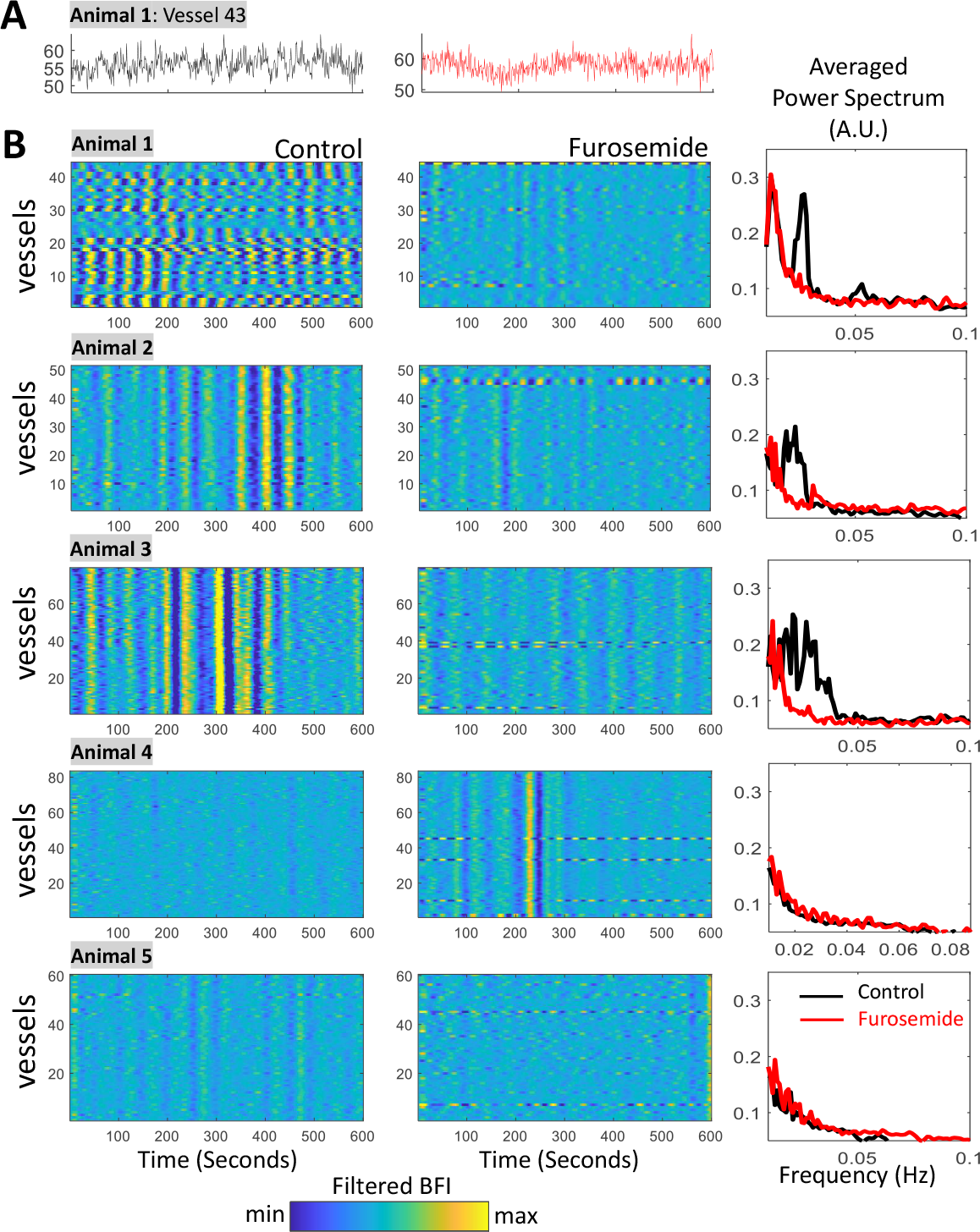
Elimination of TGF with furosemide administration in normotensive Sprague Dawley rats. A) An example time series taken from one vessel shows that the TGF oscillations present during the control period (black) disappear after furosemide infusion (red). B) Carpet plots show synchronization of TGF across many vessels with the color range representing filtered blood flow index. The first column (control) demonstrates TGF-driven oscillatory striations over the observation period. The second column (furosemide) shows vanishing TGF oscillations, demonstrated by the reduced striation compared to the first column. Third column: Average power spectra show that TGF oscillations are eliminated by furosemide (red) in all five animals.

To quantify the visual changes shown in Fig. 4, we selected three metrics to compare between control and furosemide as described in the methods: mean blood flow index (BFI), the standard deviation of the TGF bandpass filtered signal (Sigma), and the area under the curve of the power spectrum (AUC) (Fig. 5A, B, and C). When the TGF was suppressed by administrating furosemide, the BFI averaged over the observation period increased significantly (*P* < 0.0001), the Signa decreased significantly (*P* < 0.00005), and the AUC decreased significantly (*P* < 0.00005). A decrease in Sigma indicates the lack of TGF-induced oscillations in the filtered signal. The reduced AUC shows that the significance of the TGF frequency band in the power spectrum is weak. In Fig. 5D, we used a 3-dimensional scatter plot to show the spatial separation of the three metrics between control and furosemide for every segmented microvessels (n=318). While the separation of data is more evident for animals with strong TGF oscillations in the control period (animals 1-3), the spatial separation of the data points is less visible for animals that were initially without the presence of the TGF oscillations (animals 4 and 5). Linear mixed model ANOVA revealed statistical significance for all three metrics, shown in Table 1.

**Figure 5:**
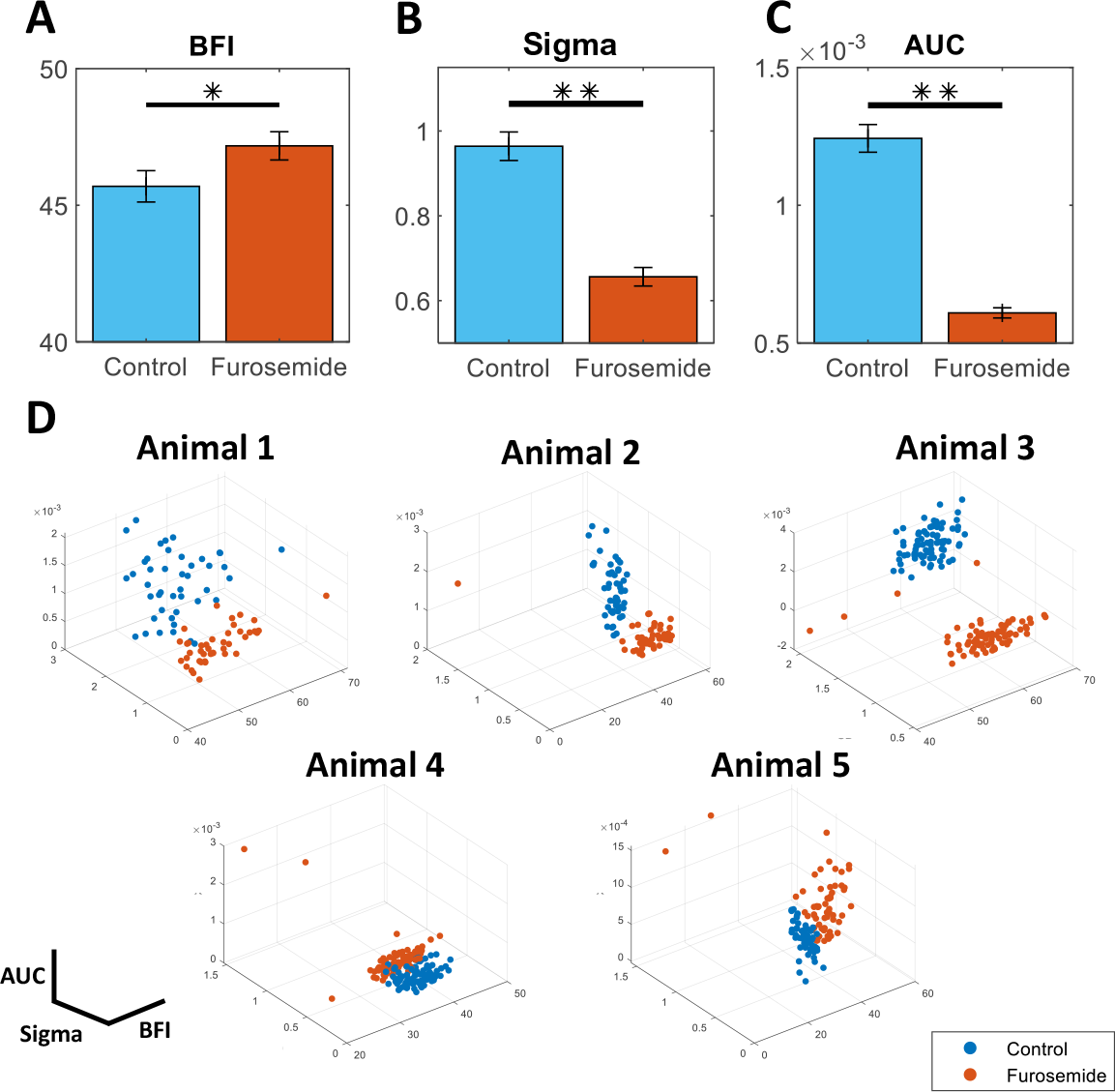
Increased local blood flow and the TGF dynamic changes induced by furosemide administration. Top panel: Bar plots represent mean ± SE for control (blue) and furosemide (orange) using three metrics: A) mean blood flow index (BFI), B) standard deviation of filtered TGF time series (Sigma), and C) area under the curve (AUC). Reduced Sigma and AUC metrics represent the elimination of TGF oscillations. D) Three metrics are plotted for every observed micro-vessel. Top row: 3 animals show a good separation in metrics between the control and the furosemide condition. Bottom row: Metrics of control and furosemide are clustered together. * corresponds to *P* < 0.0001; \*\**P* < 0.00005.

**Table 1:**
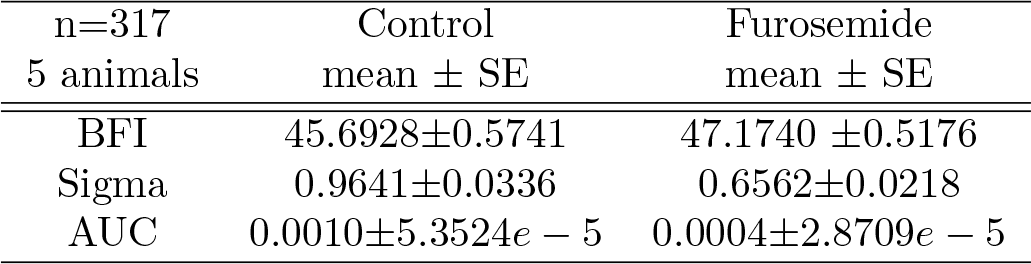
Comparison of renal hemodynamic metrics between control and furosemide.

### 3.3 Effect of acute sodium-glucose co-transporter 2 blockade on TGF dynamics

We infused phlorizin, a sodium-glucose co-transporter 2 inhibitor known to increase the urine flow and the excretion of sodium and glucose in the urine. We observed a significant increase in urine flow rate from 11.11 ±2.83 to 24.17±4.38 *μ*L/min (*P* < 0.05). The glucose excretion increased from 0.01 to 4.77 ±1.14 *μ*Eq/min (*P* < 0.05) along with increased sodium excretion from 0.56±0.27 to 3.50±0.35 *μ*Eq/min (*P* < 0.01). The blood pressure decreased from 108.29 ±5.58 to 100.73 ±4.93 mmHg, but the difference was statistically insignificant.

To explore the changes in TGF hemodynamics induced by inhibiting sodium glucose co-transporter 2, we systemically administered phlorizin in rats and observed the renal hemodynamics. As expected, Fig. 6A shows that TGF oscillations remain intact after the infusion but operate at a reduced blood flow, as exemplified on the left panel of Fig. 6A. The BFI oscillates around 60 for the control period, and the BFI oscillates around 52 for the phlorizin period. The BFI oscillation can be observed in about 40 micro-vessels in each animal, as demonstrated in Fig. 6B. In three animals (animals 1-3), TGF oscillations are evident and represented by the vertical blue-yellow striations in the control column and peaks (in black) in their respective power spectra in the third column. Animals 4, 5, and 6 lack the TGF oscillations, evidenced by the non-existent TGF frequency peak(s) in their respective power spectra. The color range does not reach the maximum yellow color intensity in the phlorizin column of Fig. 6B, which signifies a reduction in blood flow index.

**Figure 6:**
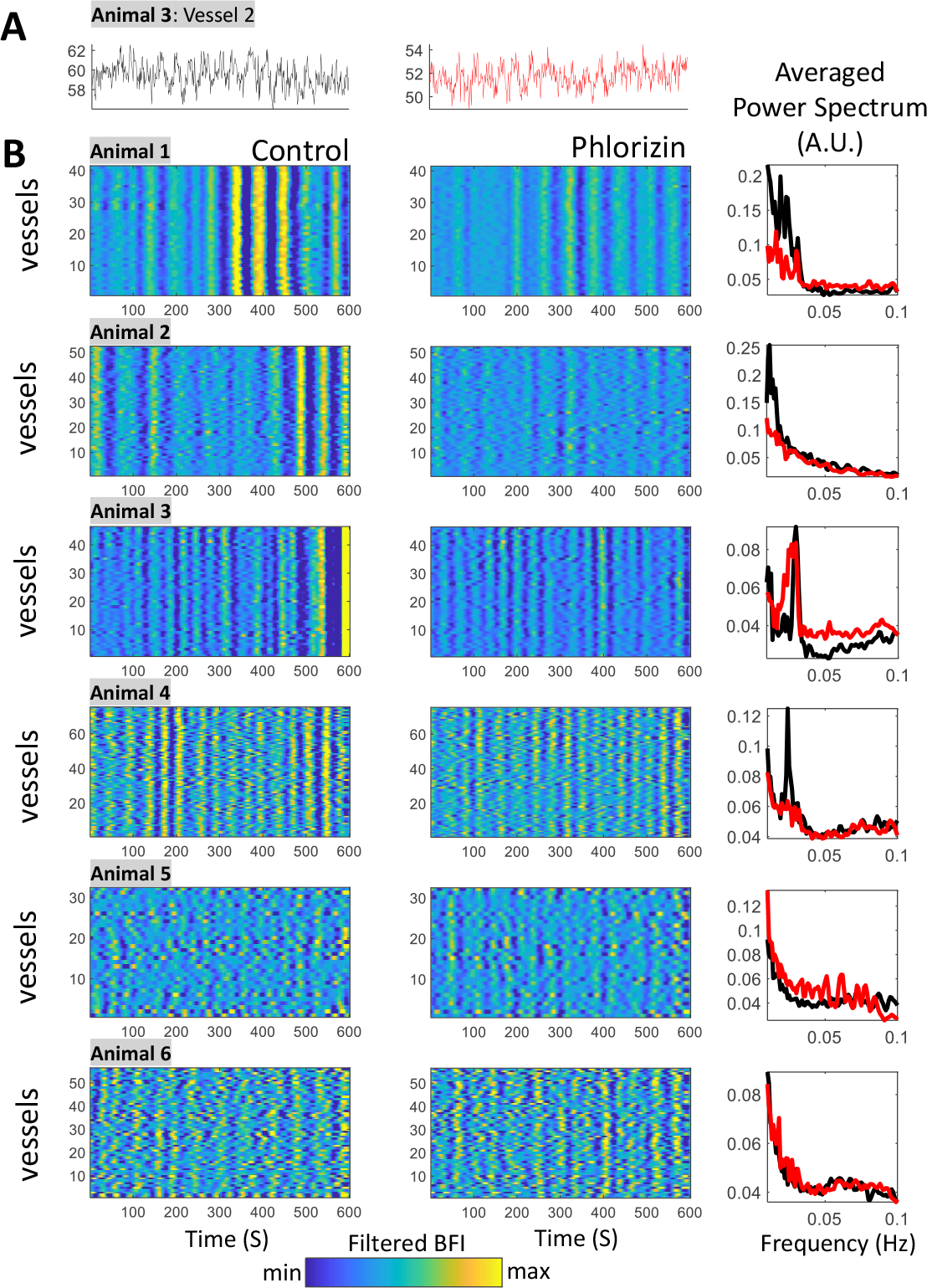
TGF in segmented vessels under phlorizin administration in normotensive Sprague Dawley rats. A) Exemplary time series of a vessel demonstrating TGF oscillations in both control and phlorizin. Please note the different y-axis B) First column (control): TGF-band filtered signals show consistent TGF oscillations over the observation time. In the second column (phlorizin): TGF oscillations are still well pronounced. Third column: Averaged power spectra show that phlorizin (red) can induce diverse responses. Animal 7 is not shown due to limited space but is included in all statistics

We quantified the same metrics for the phlorizin group as we did for the furosemide group (Fig. 5) to measure the altered TGF-mediated hemodynamics induced by phlorizin. We observed a significant decrease in blood flow (BFI) of cortical micro-vessels, indicating a reduction in glomerular perfusion across all observed nephrons (Fig. 7A). A significant decrease in Sigma indicates a smaller amplitude for TGF oscillations. The area under the curve was also reduced, indicating a reduced prominence of the TGF frequency in power spectra due to smaller amplitudes in TGF oscillations. Fig. 7D shows that the calculated metrics are well-separated between before and after the administration of phlorizin in animals 1-3 and 7, while animals 4-6 do not (animal 7 is not shown in the figure due to space). Linear mixed model ANOVA revealed that phlorizin induced a statistically significant decrease in BFI, a decrease in Sigma, and a decrease in AUC. We summarize our results in Table 2

**Figure 7:**
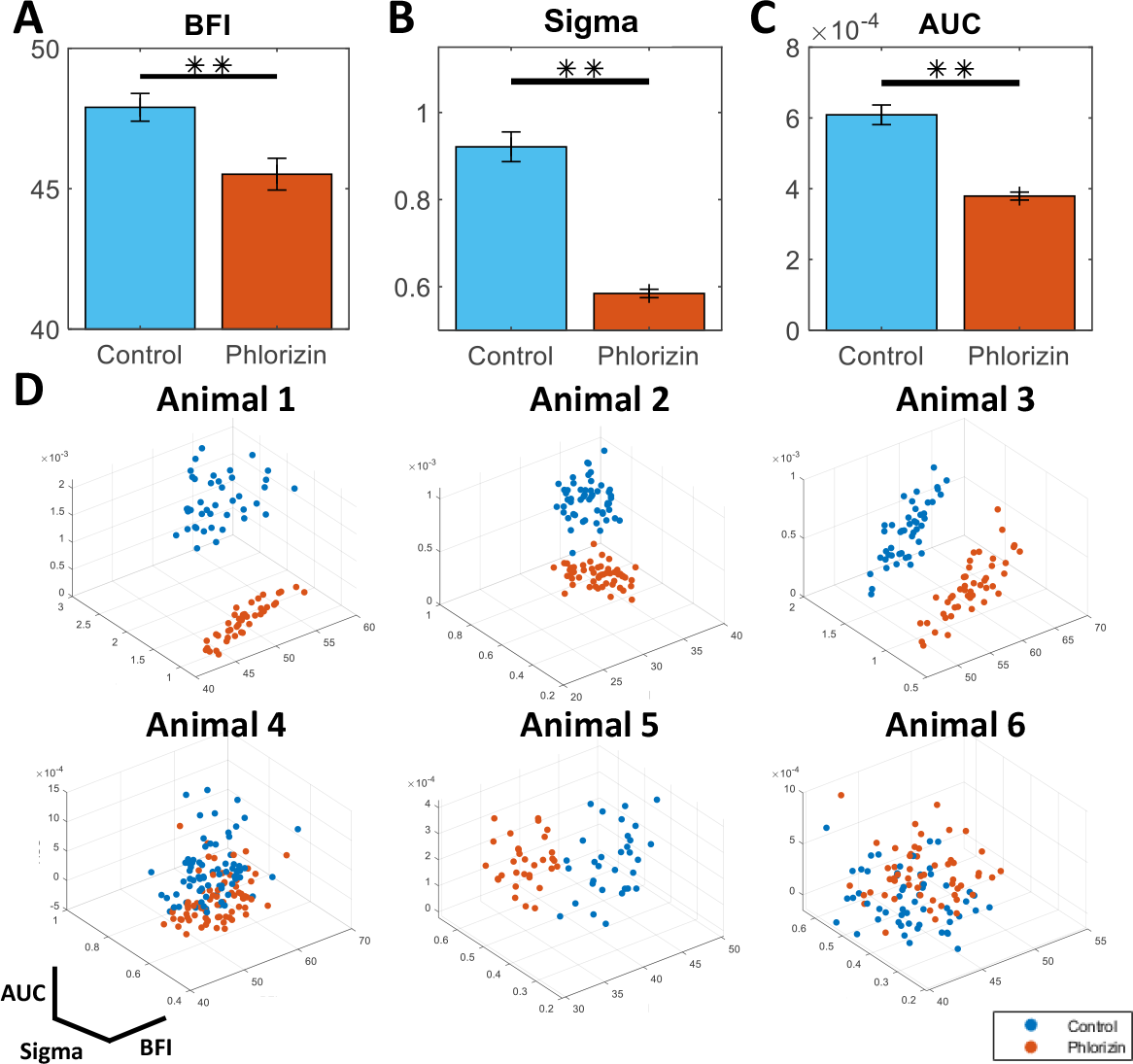
Decreased nephron blood flow and TGF dynamic changes induced with phlorizin administration. Top panel: Bar plots represent mean ± SE between control (blue) and phlorizin (orange) of three TGF metrics: (A) mean blood flow index (BFI), (B) standard deviation of filtered TGF time series (Sigma), and (C) area under the curve (AUC). D) 3-dimensional scatter plot of the three metrics for each micro-vessels (n=7, One animal was omitted for space but is included in all statistics). D) Top row: 3 animals show a good spatial separation across all three dimensions. Bottom row: control and phlorizin clusters show similar metrics in 2 animals. ** corresponds to *P* < 0.00005.

**Table 2:** Comparison of renal hemodynamic metrics between control and phlorizin.

| n=318<br>7 animals | Control<br>mean ± SE | Phlorizin<br>mean ± SE |
| --- | --- | --- |
| BFI | 47.8998 ± 0.4945 | 45.5179 ± 0.5658 |
| Sigma | 0.9213 ± 0.0341 | 0.5845 ± 0.0096 |
| AUC | 5.1796e-4 ± 2.9815e-5 | 3.0290e-4 ± 1.3805e-5 |

## 4 Discussion

This study reveals altered dynamics of TGF induced by NKCC2 and SGLT2 inhibition in a large population of nephrons for the first time, connecting the micro-hemodynamics embedded in a macroscopic nephron-vascular network. Many studies evaluating the effects of drugs on renal function rely on single nephron measurements (i.e., SNGFR) or whole animal measurements (i.e., GFR_inulin_).

There is a strong link between the changes in the pre-distal tubular reabsorption of solutes like sodium and glucose and the sequential perturbation of the TGF mechanism, which may alter the glomerular filtration pressure. The critical action of furosemide on renal hemodynamics found in this study is that the increased blood flow through an afferent arteriole supplying a glomerulus occurs at a population level due to TGF inhibition-caused by the inhibition of chloride reabsorption in the loop of Henle and the detection of chloride in the macula densa. This paper’s results are consistent with a single nephron study, which found that an intra-luminal microperfusion of furosemide can abolish TGF oscillations in Sprague-Dawley rats [18, 19].

Phlorizin, a sodium-glucose 2 inhibitor, follows a more complex narrative than furosemide as it targets the proximal tubular sodium reabsorption but not the sensing mechanism at the macula densa, i.e., TGF remains intact. Our findings show a reduction in blood flow to the nephron and reduced amplitude of TGF oscillations in response to systemic infusion of phlorizin, which is in line with several studies showing nephroprotective effects of SGLT2 inhibitors [10, 20, 11, 21]. The protective mechanism is believed to come from a decrease in glomerular capillary pressure caused by the afferent arteriolar constriction elicited by SGLT2’s effect on TGF. The TGF operating point normally resides near the midpoint of the TGF curve [22]. A steady decline in proximal reabsorption saturates the TGF response and abolishes the TGF-driven autoregulation unless the TGF somehow adapts to shift the operating point back to the steep portion of the TGF curve. Our results suggest that the reduced amplitude of the TGF-mediated blood flow oscillation is related to the shift of the TGF operating point. In agreement with our data, Thomson et al. [23] showed that inhibiting proximal HCO3 reabsorption, similar to SGLT2 inhibition, can lead to a reduction in overall proximal reabsorption. Sequentially, TGF resets to a lower operating point when the proximal reabsorption is reduced [24, 23] demonstrating the adaptability of TGF in response to fluid-content changes. In support of TGF resetting following the reduction in proximal reabsorption, Kidokoro et al. [25] showed that the administration of empagliflozin (an analog of phlorizin) for 30 minutes in mice reduced the afferent arteriolar diameter *in vivo*. The significance of pharmacologically targeting various tubular transporters to alter the nephron-vascular hemodynamics in renal pathophysiologies relating to hypertension and diabetes is highlighted in these studies. But our knowledge about the importance of TGF is limited. For example, what is its role in the short-term and long-term adaptation of salt-water balance and blood pressure regulation? Furthermore, how does impaired TGF contribute to the development of pathological conditions?

The current paradigm considers TGF as a single nephron event. The micropuncture technique remains a gold standard for answering questions in renal physiology and pathophysiology. Micropuncture methods can study TGF with single nephron resolution, establishing a relationship between tubular NaCl load and SNGFR of the same nephron [26]. However, it is known that the dynamics of neighboring glomeruli can become entrained, i.e., nephrons interact via vascular propagated signals. Our understanding of these interactions is limited.

Here, we propose to look at TGF-induced cortical hemodynamic responses through high-resolution laser speckle contrast imaging. This technique allows us to simultaneously access the TGF-mediated vascular responses in many vessels on the kidney cortex. Based on time-frequency blood flow analysis, we show how the TGF becomes suppressed with the blockage of Na-K-2Cl co-transporters and increases local microcirculatory blood flow by 11.3%. However, the administration of an SGLT2 inhibitor gives the opposite results. Our metrics associated with the amplitude (Sigma) and the significance (AUC) of TGF oscillations show a significant decrease in TGF-mediated hemodynamic responses, pointing to a shift in the TGF operating point. Different animals exhibit various dynamical patterns (Fig. 6). We see a relative drop in local microcirculatory blood flow in the whole field of view by 6.42%, compared to baseline. The magnitude of change can be related to the strength of vascular responses and how many vessels respond. When we visualize TGF oscillations of phlorizin and furosemide, in Fig. 6 and Fig. 5 respectively, one can see that the TGF oscillations are preserved for the phlorizin group. In diabetes, there is an increased expression and activity of SGLT2, leading to a higher potency of SGLT2 inhibitors [21], and our approach might reveal unexpected adaptive patterns in normotensive rats.

The afferent arteriolar tone is controlled by pressure-induced (myogenic) vasomotion and by the TGF. These mechanisms contribute to efficient autoregulation [2, 27, 28]. Yip et al. [28] showed that there are two oscillating components in spontaneously fluctuating single-nephron blood flow obtained from Sprague-Dawley rats: a slow oscillation (20–30 mHz) that is mediated by TGF and a fast oscillation (100 mHz) that is related to the myogenic activity. It was shown [29] that TGF modulated myogenic activity. Yip et al. [30] observed that myogenic oscillations are enhanced when TGF was inhibited with furosemide. This study focused on TGF responses, but future studies should also consider the myogenic response. It would be interesting to see if the myogenic mechanism can compensate for the reduced TGF oscillations observed after inhibition of SGLT2. The ramifications of reduced autoregulatory efficiency are unclear: Whether this is a sign of renal decline or a mechanism used by the kidney to respond to a physiological imbalance is unclear.

In previous studies of TGF dynamics, the TGF signal was designated by choosing a single peak between a tight frequency range (0.018 Hz-0.033 Hz). However, applying time-frequency superlet analysis to renal hemodynamics unveils qualities of TGF that were neglected with previous time frequency analyses such as Fourier and Wavelet analyses. For example, fig 3 shows different kinds of time-frequency TGF responses from individual vessels in the cortex of the kidney. A single sharp peak in the power spectrum indicates stable TGF oscillations over the observation period. But multiple peaks can be attributed to (i) vascular signals originating from different nephrons that operate at different TGF frequencies or (ii) a change in the TGF frequency of a single nephron over an observation period in response to an external perturbation. However, a detailed analysis of this would require a combination of high-resolution blood flow imaging and structural imaging techniques.

Network behavior forms an integral part of renal autoregulation. TGF refers to the feedback regulation of glomerular filtration rate in a single nephron based on sensory information about the distal tubule fluid. Recent experiments showed that many nephrons coordinate their TGF-induced hemodynamic responses. The presence of TGF oscillations links to the synchronization of vascular responses of neighboring nephrons [31, 32, 15]. Recently, Postnov et al. [13] confirmed that blood flow in renal microcirculation tends to demonstrate clustered, frequency-locked activity. Groups of vessels exhibit a short or long-term synchronous behavior of TGF oscillations that is disengaged once TGF is eliminated with furosemide (Fig. 4). One could expect that renal autoregulation provides better protection when nephrons act together. Cooperating nephrons can increase the efficiency of renal autoregulation by collectively engaging in more preglomerular resistance. Yet, the efficacy of such cooperative behavior on overall renal autoregulation remains an open question.

## 5 Acknowledgment

Karin Larsen assisted in animal preparations. This work is supported by the Novo Nordisk Foundation (Grant NNF18OC0052728).

## 6 Data availability

Data underlying the results presented in this paper are not publicly available but may be obtained from the authors upon reasonable request.

